# A learned embedding for efficient joint analysis of millions of mass spectra

**DOI:** 10.1101/483263

**Authors:** Wout Bittremieux, Damon H. May, Jeffrey Bilmes, William Stafford Noble

## Abstract

Computational methods that aim to exploit publicly available mass spectrometry repositories primarily rely on unsupervised clustering of spectra. Here, we propose to train a deep neural network in a supervised fashion based on previous assignments of peptides to spectra. The network, called “GLEAMS,” learns to embed spectra into a low-dimensional space in which spectra generated by the same peptide are close to one another. We use GLEAMS as the basis for a large-scale spectrum clustering, detecting groups of unidentified, proximal spectra representing the same peptide, and we show how to use these clusters to explore the dark proteome of repeatedly observed yet consistently unidentified mass spectra. We provide a software implementation of our approach, along with a tool to quickly embed additional spectra using a pre-trained model, to facilitate large-scale analyses.

---

In proteomics, the dominant approach to assigning peptide sequences to tandem mass spectrometry (MS/MS) data is to treat each spectrum as an independent observation during sequence database searching.^1^ However, as public data repositories have grown to include billions of MS/MS spectra over the last decade,^2^ efforts have been undertaken to make the spectra useful to researchers analyzing new datasets. For example, spectrum clustering can be used to filter for high-quality MS/MS spectra that are repeatedly observed across multiple datasets.^3–6^ Standard clustering is problematic, however, because it is an *unsupervised* approach. The input to a clustering algorithm is an unlabeled set of spectra. In practice, the labels (i.e. the associated peptide sequences) are used only in a *post hoc* fashion, to choose how many clusters to produce or to split up large clusters associated with multiple peptides.

In recent years, a revolution has occurred in machine learning, with deep neural networks proving to have applicability across a wide array of problems.^7^ Accordingly, within the field of proteomics, deep neural networks have been applied to several problems, including *de novo* peptide sequencing^8,9^ and simulating MS/MS spectra.^10,11^ However, to our knowledge no one has yet applied deep neural networks to the problem of making public repository data contribute to the analysis of new mass spectrometry experiments. We hypothesize that we can obtain more accurate and useful information about a large collection of spectra by using a supervised deep learning method that directly exploits peptide-spectrum assignments during joint analysis. Specifically, we posit that peptide labels can be used during training of a large-scale learned model of MS/MS spectra to achieve a robust, efficient, and accurate model.

Accordingly, we propose GLEAMS (GLEAMS is a Learned Embedding for Annotating Mass Spectra), which is a deep neural network that has been trained to embed MS/MS spectra into a 32-dimensional space in such a way that spectra generated by the same peptide, with the same post-translational modifications (PTMs) and charge, are close together. The learned spectrum embedding offers the advantage that new spectra can efficiently be mapped to the embedded space without requiring re-training. Our approach is fundamentally different from previous (unsupervised) spectrum clustering applications, in the sense that it uses peptide assignments generated from database search methods as labels in a supervised learning setting.

GLEAMS consists of two identical instances of an embedding neural network in a Siamese network set-up^12^ (Figure 1A). During training, the network receives pairs of spectra as input and labels indicating whether the spectra correspond to the same peptide sequence or not. Each input spectrum is encoded using three sets of features representing attributes of their precursor ion, binned fragment intensities, and similarities to an invariant set of reference spectra. Each of the different feature types is processed through a separate deep neural subnetwork, after which the outputs of the three networks are concatenated and passed to a final, fully-connected layer to produce vector embeddings with dimension 32 (Supplementary Figure 1). The entire network is trained to transform the input spectra into 32-dimensional embeddings by optimizing a contrastive loss function.^12^ Intuitively, this loss function “pulls” the embeddings of spectra corresponding to the same peptide together, and “pushes” the embeddings of spectra corresponding to different peptides apart. The embedder network thus constitutes a function that transforms spectra into latent embeddings so that spectra corresponding to the same peptide are close to each other.

**Figure 1:**
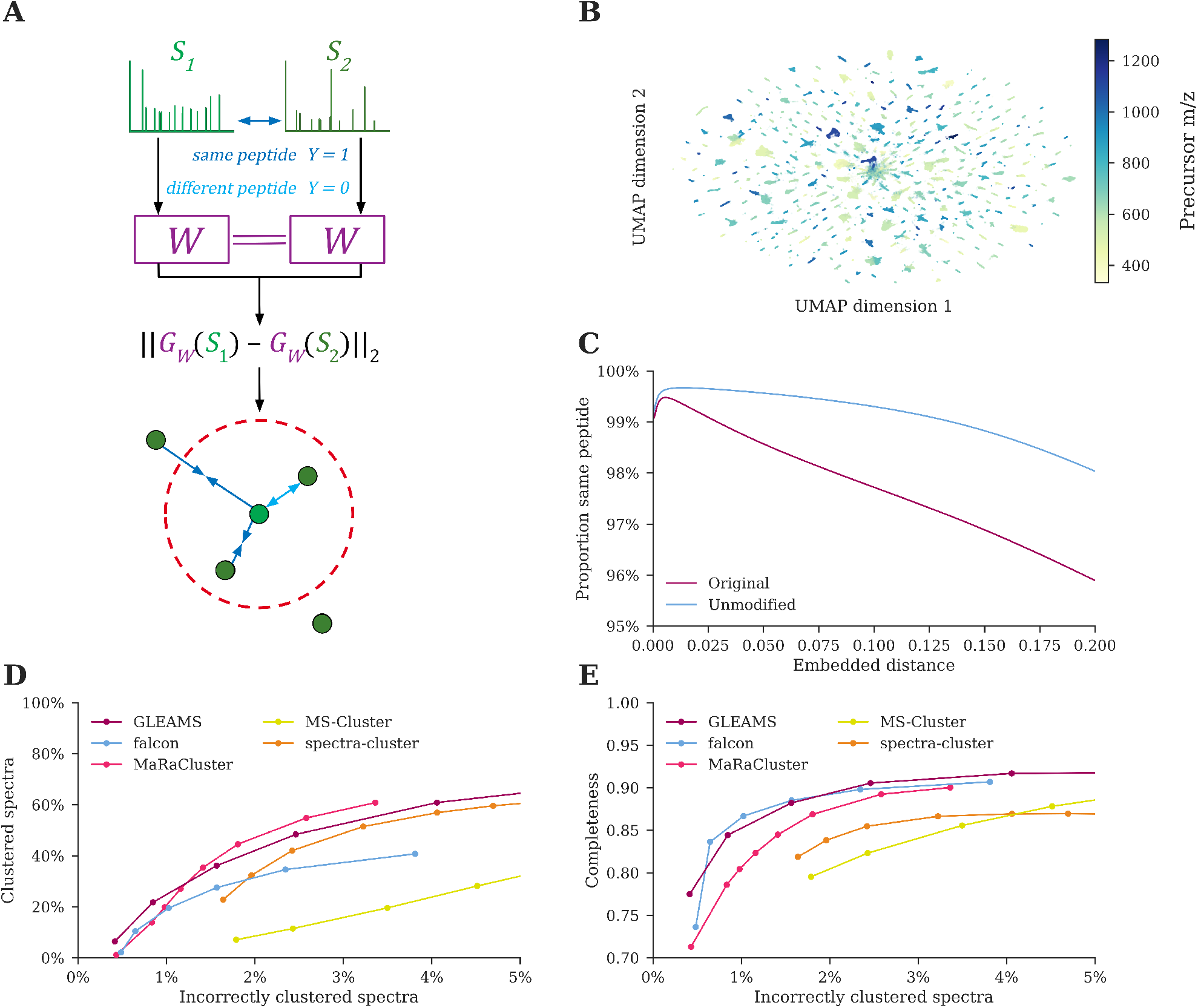
**(A)** Two spectra, *S*_1_ and *S*_2_, are encoded to vectors and passed as input to two instances of the embedder network with tied weights. The Euclidean distance between the two resulting embeddings, *G_W_* (*S*_1_) and *G_W_* (*S*_2_), is passed to a contrastive loss function that penalizes dissimilar embeddings that correspond to the same peptide and similar embeddings that correspond to different peptides, up to a margin of 1. **(B)** UMAP projection of 685 337 embeddings from frequently occurring peptides in 10 million randomly selected identified spectra from the test dataset. As indicated by the coloration, precursor *m/z* has a large influence on the location of spectra in the embedded space. Note that the visualization may group peptides with similarities on some dimensions of the 32-dimensional embedding space but which are nevertheless distinguishable based on their full embeddings. **(C)** Proportion of neighbors that have the same peptide label as a function of the distance threshold for 186 865 330 pairwise distances between 10 million randomly selected embeddings from the test dataset. Embeddings at small distances represent the same peptide (“Original”), while the majority of close neighbors with different peptide labels correspond to peptides with ambiguously localized modifications (“Unmodified”). **(D+E)** Average clustering performance over three random folds of the test dataset containing 28 million MS/MS spectra each. **(D)** The number of clustered spectra versus the number of incorrectly clustered spectra per clustering algorithm. GLEAMS and MaRaCluster succeed in clustering the highest number of spectra. MS-Cluster and spectra-cluster are unable to produce highly pure clusters, even with the most stringent hyperparameters. **(E)** Cluster completeness versus the number of incorrectly clustered spectra per clustering algorithm. GLEAMS and falcon produce the most complete clustering results. This indicates that GLEAMS achieves a greater data reduction than alternative clustering tools, without sacrificing the cluster quality.

GLEAMS was trained using a set of 30 million high-quality peptide-spectrum matches (PSMs) derived from the MassIVE knowledge base (MassIVE-KB).^6^ Importantly, peptide sequence information is only required during initial supervised training of the Siamese network. Subsequent processing using an individual embedder instance is agnostic to the peptide labels and can be performed on identified and unidentified spectra in a similar fashion. After training, the embedder model was used to process 669 million spectra from 227 public human proteomics datasets included in MassIVE-KB. As an initial evaluation of the learned embeddings, these spectra were further projected down to two dimensions using UMAP^13^ for visual inspection. The visualizations suggest that precursor mass (Figure 1B) and precursor charge (Supplementary Figure 2) strongly influence the structure of the embedded space, and that similar spectra are indeed located close to each other. Additionally, several of the individual embedding dimensions show a correlation with the precursor mass, peptide sequence length, or whether the peptides have an arginine or lysine terminus (Supplementary Table 1). This indicates that the GLEAMS embeddings capture latent characteristics of the spectra. Interestingly, although some of these properties were provided as input to the neural network, such as precursor mass, other properties were derived from the data without explicitly encoding them.

If our training worked well, then spectra generated by the same peptide should lie close together, according to a Euclidean metric, in the embedded space. Accordingly, we investigated, for 10 million randomly chosen embedded spectra, the relationship between neighbor distance and the proportion of labeled neighbors that have the same peptide label. The results show that neighbors at small distances overwhelmingly represent the same peptide (Figure 1C). Furthermore, the few different-peptide labels at very small distances almost entirely represent virtually indistinguishable spectra that have identical peptide labels but differ in ambiguous modification localizations. We also investigated the false negative rate, for 10 million randomly chosen embedding pairs, to understand the extent to which embeddings that correspond to the same peptide are distant in the embedded space (Supplementary Figure 3). This analysis shows an excellent separation between same-labeled embeddings and embeddings corresponding to different peptides, with a very small false negative rate of only 1% at a distance threshold corresponding to 1% false discovery rate (FDR). Furthermore, the embeddings are robust to different types of mass spectrometry data. Phosphorylation modifications were not included in the MassIVE-KB dataset, and GLEAMS thus did not see any phosphorylated spectra during its training. Nonetheless, GLEAMS was able to embed spectra from a phosphoproteomics study with high accuracy (Supplementary Figure 4).^14^

To further investigate the utility of the GLEAMS embedding, we performed clustering in the embedded space to find groups of similar spectra, and we compared the performance to that of the spectrum clustering tools MS-Cluster,^3^ spectra-cluster,^4,5^ MaRaCluster,^15^ and falcon^16^. The comparison indicates that clustering in the GLEAMS embedded space is of a similar or higher quality than clusterings produced by state-of-the-art tools (Figure 1D–E). Additionally, GLEAMS generates highly “complete” clustering results (Figure 1E). Completeness measures the extent to which multiple spectra corresponding to the same peptide are concentrated in few clusters. Compared to alternative clustering tools, GLEAMS produces larger clusters, with data drawn from more diverse studies (Supplementary Figure 5). By minimizing the extent to which spectra generated from the same peptide are assigned to different clusters, GLEAMS achieves improved data reduction from spectrum clustering compared to alternative clustering tools. Furthermore, GLEAMS achieves excellent performance irrespective of the clustering algorithm used (Supplementary Figure 6). This indicates that, despite their compact size, the GLEAMS embeddings are rich in information and suitable for downstream processing. We hypothesize that GLEAMS’ supervised training allows the model to focus on relevant spectrum features while ignoring confounding features—for example, peaks corresponding to a ubiquitous contaminant within a single study, boosting intra-study spectrum similarity. This property is especially relevant when performing spectrum clustering at the repository scale, to maximally reduce the volume of heterogeneous data for efficient downstream processing.

A key outstanding question in protein mass spectrometry analysis concerns the source of spectral “dark matter,” i.e. spectra that are observed repeatedly across many experiments but consistently remain unidentified. Frank et al. [17] have previously used MS-Cluster to identify 4 million unknown spectra included in “spectral archives,” and Griss et al. [5] have used spectra-cluster to obtain identifications for 9 million previously unannotated spectra in the PRoteomics IDEntifications (PRIDE) repository. The original MassIVE-KB results^6^ include identifications for 185 million PSMs out of 669 million MS/MS spectra (1% FDR), leaving a vast amount of spectral data unexplored.

To characterize the unidentified spectra, we performed GLEAMS clustering (~1% incorrectly clustered spectra) to group 511 million spectra in 60 million clusters, followed by a multi-step procedure to explore the dark proteome. This procedure involved propagating peptide labels within clusters, as well as targeted open modification searching of representative spectra drawn from clusters of unidentified spectra (see Methods). In total, this strategy succeeded in assigning peptides to 132 million previously unidentified PSMs, increasing the number of identified spectra by 71% (Figure 2A). Additionally, there are 207 million clustered spectra that remained unidentified. Because these spectra are repeatedly observed and expected to be high-quality, they likely correspond to true signals. Consequently, this is an important collection of spectra to investigate using newly developed computational methods to further explore the dark proteome. The open modification searching results also provided information on the presence of PTMs in the human proteome (Figure 2B, Supplementary Data 1). Besides abundant modifications that can be artificially introduced during sample processing, such as carbamidomethylation and oxidation, biologically relevant modifications from enrichment studies, such as phosphorylation, were frequently observed. We provide all of these data as a valuable community resource to further explore the dark proteome.

**Figure 2:**
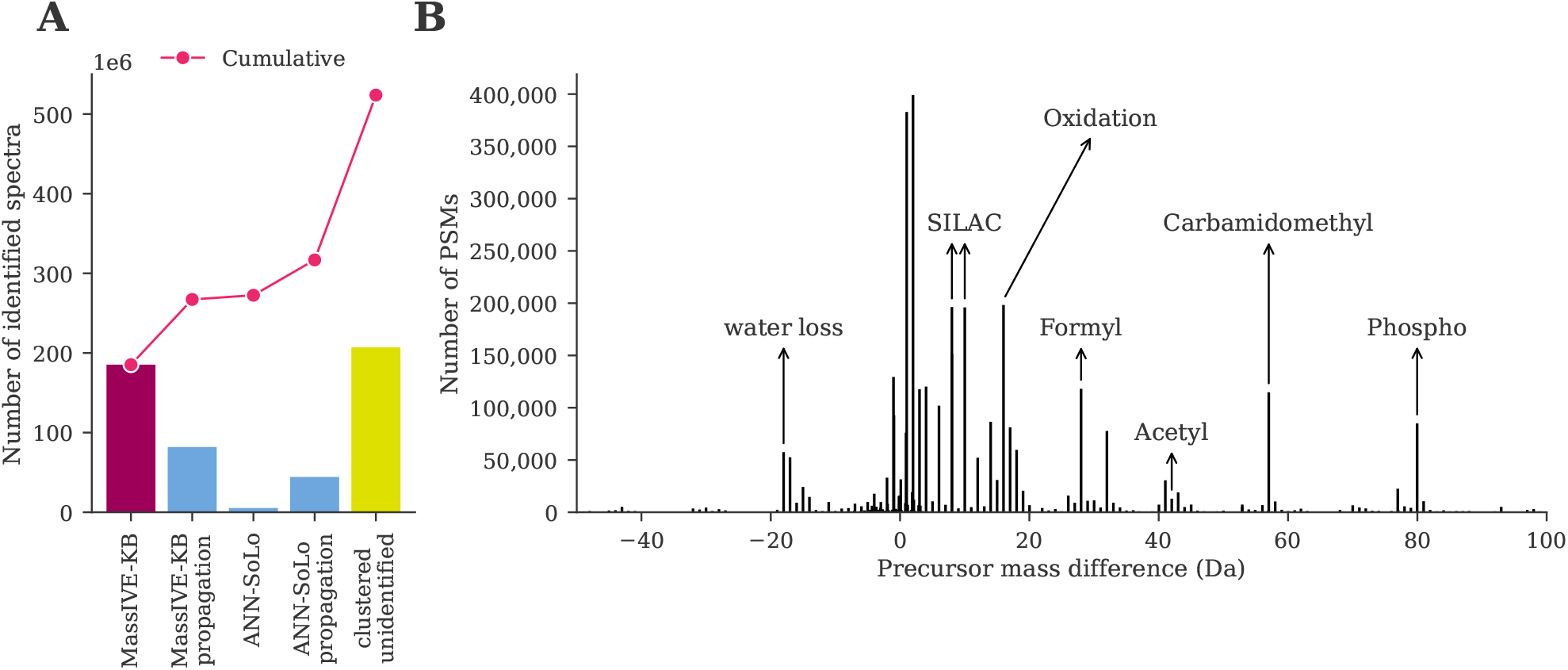
Exploration of the dark proteome using GLEAMS to process previously unidentified spectra. **(A)** GLEAMS succeeded in identifying 71% additional PSMs (blue) compared to the original MassIVE-KB results (dark pink) by performing targeted open modification searching of cluster medoid spectra and propagating peptide labels within clusters. Several high-quality clustered, yet unidentified spectra (yellow) remain to further explore the dark proteome. **(B)** Precursor delta masses observed from open modification searching. Some of the most frequent delta masses are annotated with their likely modifications, sourced from Unimod.^18^ See Supplementary Table 2 for details of the top 500 observed precursor mass differences.

We have demonstrated the utility of the 32-dimensional embedding learned by GLEAMS. By mapping spectra from diverse experiments into a common latent space, we can efficiently add an additional 71% to the identifications derived from database search. A key factor in GLEAMS’ strong performance is its unique ability to efficiently operate on hundreds of millions to billions of spectra, corresponding to the size of an entire proteomics repository (Supplementary Figure 7). Once the embedder is trained, new spectra representing previously unobserved peptides can be embedded and used for analysis without performing any expensive operations as long as they have sufficiently similar characteristics to the distribution of training spectra. This makes it possible in principle to assign new spectra to spectrum clusters nearly instantaneously upon submission to a repository, giving researchers the immediate benefit of the combined analysis efforts of the entire proteomics community.

## Methods

### Encoding mass spectra for network input

Each spectrum is encoded as a vector of 3010 features of three types: precursor attributes, binned fragment intensities, and dot product similarities with a set of reference spectra.

Precursor mass, *m/z*, and charge are encoded as a combined 61 features. Precursor mass and *m/z* are each extremely important values for which precision is critical, and so they are poorly suited for encoding as single input features for a neural network. Accordingly, we experimented with several binary encodings of precursor mass and *m/z*, each of which gave superior performance on validation data than a real-value encoding, and settled on the encoding that gave moderately better performance than the others: a 27-bit “Gray code” binary encoding, in which successive values differ by only a single bit, preserving locality and eliminating thresholds at which many bits are flipped at once. Precursor values may span the range 400 Da to 6000 Da, so the Gray code encoding has a resolution of 4 × 10^-5^ Da. Fragment values may span the range 50.5*m/z* to 2500*m/z*, so the Gray code encoding has a resolution of 2 × 10^-5^*m/z*. Spectrum charge is one-hot encoded: seven features represent charge states 1–7, all of which are set to 0 except the charge corresponding to the spectrum (spectra with charge 8 or higher are encoded as charge 7).

Fragment peaks are encoded as 2449 features. Fragment intensities are square-root transformed and then normalized by dividing by the sum of the square-root intensities. Fragments outside the range 50.5*m/z* to 2500*m/z* are discarded, and the remaining fragments are binned into 2449 bins at 1.000 507 9*m/z*, corresponding to the distance between the centers of two adjacent clusters of physically possible peptide masses,^19^ with bins offset by half a mass cluster separation width so that bin boundaries fall between peaks. This bin size was chosen in order to accommodate data acquired using various instruments and protocols, and in deference to practical constraints on the number of input features for the deep learning approach. Consequently, it is unrelated to the optimal fragment mass tolerance for database search for a given run.

Similarities of each spectrum to an invariant set of reference spectra are encoded as 500 features. Each such feature is the normalized dot product between the given spectrum and one of an invariant set of 500 reference spectra chosen randomly from the training dataset. This can be considered as an “empirical kernel map,”^20^ allowing GLEAMS to represent the similarity between two spectra *A* and *B* by “paths” of similarities through each one of the reference spectra *R* via the transitive property; i.e., *A* is similar to *B* if *A* is similar to *R* and *R* is similar to *B*. In contrast to the fragment binning strategy described previously, similarities to the reference spectra are computed at native resolution. The 500 reference MS/MS spectra were selected from the training data by using submodular selection, as implemented in the apricot Python package (version 0.4.1).^21^ First, 1000 peak files were randomly selected from the training data, containing 22 million MS/MS spectra that were downsampled to 200 000 MS/MS spectra. These spectra were used to compute a pairwise similarity matrix (normalized dot product with fragment *m/z* tolerance 0.05*m/z*) that was used to perform submodular selection using the facility location function to select 500 representative reference spectra (Supplementary Figure 8).

Ablation tests show that optimal training performance is reached when the neural network receives all three features types as input (Supplementary Figure 9). These empirical results indicate that each of the features provides complementary information to the neural network.

### Repository-scale MS/MS data

A large-scale, heterogeneous dataset derived from the MassIVE knowledge base (MassIVE-KB; version 2018-06-15)^6^ was used to develop GLEAMS. As per Wang et al. [6], the MassIVE-KB dataset consists of 31 TB of human data from 227 public proteomics datasets. In total 28 155 peak files in the mzML^22^ or mzXML format were downloaded from MassIVE, containing over 669 million MS/MS spectra.

All spectra were processed using a uniform identification pipeline during initial compilation of the MassIVE-KB dataset.^6^ MSGF+^23^ was used to search the spectra against the UniProt human reference proteome database (version May 23, 2016).^24^ Cysteine carbamidomethylation was set as a fixed modification, and variable modifications were methionine oxidation, N-terminal acetylation, N-terminal carbamylation, pyroglutamate formation from glutamine, and deamidation of asparagine and glutamine. MSGF+ was configured to allow one ^13^C precursor mass isotope, at most one non-tryptic terminus, and 10 ppm precursor mass tolerance. The searches were individually filtered at 1% PSM-level FDR. A dynamic search space adjustment was performed during processing of the synthetic peptide spectra from the ProteomeTools project^25^ and affinity purification mass spectrometry runs from the BioPlex project^26^ to account for differences in sample complexity and spectral characteristics.^6^ Next, the MassIVE-KB spectral library was generated using the top 100 PSMs for each unique precursor (i.e. combination of peptide sequence and charge), corresponding to 30 million high-quality PSMs (uniformly 0% PSM-level FDR from the original searches).^6^

The MSGF+ identification results for the full MassIVE-KB dataset were obtained from MassIVE in the mzTab format^27^ and combined in a single metadata file containing 185 million PSMs. Additionally, information for the 30 million filtered PSMs to create the MassIVE-KB spectral library was independently retrieved.

### Neural network architecture

The embedder network (Supplementary Figure 1) takes each of the three types of inputs separately. The precursor features are processed through a two-layer fully-connected network with layer dimensions 32 and 5. The 2449-dimensional binned fragment intensities are processed through five blocks of one-dimensional convolutional layers and max pooling layers, inspired by the VGG architecture.^28^ The first two blocks consist of two consecutive convolutional layers, followed by a max pooling layer. The third, fourth, and fifth blocks each consist of three consecutive convolutional layers, followed by a max pooling layer. The number of output filters of each of the convolutional layers is 30 for the first block, 60 for the second block, 120 for the third block, and 240 for the fourth and fifth blocks. All blocks use convolutional layers with convolution window length 3 and convolution stride length 1. All max pooling layers consist of pool size 1 and stride length 2. In this fashion, the first dimension is halved after every block to ultimately convert the 2449×1 dimensional input tensor to a 71×240 dimensional output tensor. The 500-dimensional reference spectra features are processed through a two-layer fully-connected network with layer dimensions 750 and 250. The output of the three networks is concatenated and passed to a final, L2-regularized, fully-connected layer with dimension 32.

All network layers use the scaled exponential linear units (SELU) activation function.^29^ The fully-connected layers are initialized using LeCun normal initialization,^30^ and the convolutional layers are initialized using the Glorot uniform initialization.^31^

To train the embedder, we construct a “Siamese network” containing two instances of the embedder with tied weights *W* forming function *G_W_* (Figure 1A). Pairs of spectra *S*_1_ and *S*_2_ are transformed to embeddings *G_W_*(*S*_1_) and *G_W_*(*S*_2_) in each instance of the Siamese network, respectively. The output of the Siamese network is the Euclidean distance between the two embeddings: ||*G_W_*(*S*_1_) – *G_W_*(*S*_2_)||_2_. The Siamese network is trained to optimize the following contrastive loss function:^12^

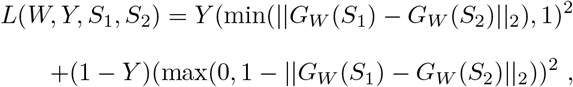

where *Y* is the label associated with the pair of spectra *S*_1_ and *S*_2_.

### Training the embedder

The GLEAMS model was trained using the 30 million high-quality PSMs used for compilation of the MassIVE-KB spectral library. PSMs were randomly split by their MassIVE dataset identifier so that the training, validation, and test sets consisted of approximately 80%, 10%, and 10% of all PSMs respectively (training set: 24 986 744 PSMs / 554 290 510 MS/MS spectra from 184 datasets; validation set: 2 762 210 PSMs / 30 386 035 MS/MS spectra from 11 datasets; test set: 2 758 019 PSMs / 84 699 214 MS/MS spectra from 24 datasets).

The Siamese neural network was trained using positive and negative spectra pairs. Positive pairs consist of two spectra with identical precursors, and negative spectra consist of two spectra that correspond to different peptides within a 10 ppm precursor mass tolerance with at most 25% overlap between their theoretical b and y fragments. In total 317 million, 205 million, 43 million, and 5 million positive training pairs were generated for precursor charges 2 to 5, respectively; and 8.347 billion, 3.263 billion, 182 million, and 5 million negative training pairs were generated for precursor charges 2 to 5, respectively.

The Siamese neural network was trained for 50 iterations using the rectified Adam optimizer^32^ with learning rate 0.0002. Each iteration consisted of 40 000 steps with batch size 256. The pair generators per precursor charge and label (positive/negative) were separately shuffled and rotated to ensure that each batch consisted of an equal number of positive and negative pairs and balanced precursor charge states. After each iteration the performance of the network was assessed using a fixed validation set consisting of up to 512 000 spectrum pairs per precursor charge.

Training and evaluation were performed on a Intel Xeon Gold 6148 processor (2.4 GHz, 40 cores) with 768 GB memory and four NVIDIA GeForce RTX 2080 Ti graphics cards.

### Phosphoproteomics embedding

An independent phosphoproteomics dataset by Hijazi et al. [14], generated to study kinase network topology, was used to evaluate the robustness of the GLEAMS embeddings for unseen post-translational modifications. All raw and mzIdentML^33^ files were downloaded from PRIDE (project PXD015943) using ppx (version 1.1.1)^34^ and converted to mzML files^22^ using ThermoRawFileParser (version 1.3.4).^35^ As per Hijazi et al. [14], the original identifications were obtained by searching with Mascot (version 2.5)^36^ against the SwissProt database (SwissProt_Sep2014_2015_12.fasta), with search settings of up to two tryptic missed cleavages; precursor mass tolerance 10 ppm; fragment mass tolerance 0.025 Da; cysteine carbamidomethylation as a fixed modification; and N-terminal pyroglutamate formation from glutamine, methionine oxidation, and phosphorylation of serine, threonine, and tyrosine as variable modifications. The identification results included 3.7 million PSMs at 1% FDR (of which 98.5% are phosphorylated) for 18.6 million MS/MS spectra. All spectra were embedded with the previously trained GLEAMS model, and 1.185 billion positive pairs consisting of PSMs with identical peptide sequences and 293 million negative pairs consisting of PSMs with different sequences within a 10 ppm precursor mass tolerance were generated.

### Embedding clustering

Prior to clustering, the MS/MS spectra were converted to embeddings using the trained GLEAMS model. Next, the embeddings were split per precursor charge and partitioned into buckets based on their corresponding precursor mass so that the precursor *m/z* difference of consecutive embeddings in neighboring buckets exceeded the 10 ppm precursor *m/z* tolerance. Embeddings within each bucket were clustered separately based on their Euclidean distances. Different clustering algorithms were used, including hierarchical clustering with complete linkage, single linkage, and average linkage, and DBSCAN clustering.^37^ An important advantage of these clustering algorithms is that the number of clusters is not required to be known in advance.

In some cases, jointly clustered embeddings would violate the 10 ppm precursor mass tolerance because embeddings within a cluster were connected through other embeddings with intermediate precursor mass. To avoid such false positives, the clusters were postprocessed by hierarchical clustering with complete linkage of the cluster members’ precursor masses. In this fashion, clusters were split into smaller, coherent clusters so that none of the embeddings in a single cluster had a pairwise precursor mass difference that exceeded the precursor mass tolerance.

### Cluster evaluation

Five clustering algorithms—GLEAMS clustering, falcon^16^, MaRaCluster,^15^ MS-Cluster,^3^ and spectra-cluster^4,5^—were run using a variety of parameter settings for each. For GLEAMS clustering, several clustering algorithms were used. For hierarchical clustering with complete linkage, Euclidean distance thresholds of 0.1, 0.2, 0.3, 0.4, 0.5, 0.6, 0.7, and 0.8 were used. For hierarchical clustering with single linkage, Euclidean distance thresholds of 0.05, 0.10, 0.15, 0.20, and 0.25 were used. For hierarchical clustering with average linkage, Euclidean distance thresholds of 0.1, 0.2, 0.3, 0.4, 0.5, and 0.6 were used. For DBSCAN clustering, Euclidean distance thresholds of 0.005, 0.01, 0.02, 0.03, 0.04, 0.05, 0.06, 0.07, 0.08, 0.09, and 0.10 were used. Falcon (version 0.1.3)^16^ was run with a precursor mass tolerance of 10 ppm, fragment mass tolerance 0.05 Da, minimum fragment intensity 0.1, and square root intensity scaling. Cosine distance thresholds were 0.01, 0.05, 0.10, 0.15, 0.20, and 0.25. Other options were kept at their default values. MaRaCluster (version 1.01)^15^ was run with a precursor mass tolerance of 10 ppm, and with identical P-value and clustering thresholds −3.0, −5.0, −10.0, −15.0, −20.0, −25.0, −30.0, or −50.0. Other options were kept at their default values. MS-Cluster (version 2.00)^3^ was run using its “LTQ_TRYP” model for three rounds of clustering with mixture probability 0.000 01, 0.0001, 0.001, 0.005, 0.01, 0.05, or 0.1. The fragment mass tolerance and precursor mass tolerance were 0.05 Da and 10 ppm, respectively, and precursor charges were read from the input files. Other options were kept at their default values. spectra-cluster (version 1.1.2)^4,5^ was run in its “fast mode” for three rounds of clustering with the final clustering threshold 0.999 99, 0.9999, 0.999, 0.99, 0.95, 0.9, or 0.8. The fragment mass tolerance and precursor mass tolerance were 0.05 Da and 10 ppm, respectively. Other options were kept at their default values.

The clustering tools were evaluated using 84 million MS/MS spectra originating from 24 datasets in the test set. The spectra were split in three randomly generated folds, containing approximately 28 million MS/MS spectra each, and exported to MGF files for processing using the different clustering tools. To evaluate cluster quality, the mean performance over the three folds was used. Valid clusters were required to consist of minimum five spectra, and the remaining spectra were considered as unclustered.

The following evaluation measures were used to assess cluster quality:

#### Clustered spectra

The number of clustered spectra divided by the total number of spectra.

#### Incorrectly clustered spectra

The number of incorrectly clustered spectra divided by the total number of clustered, identified spectra. Spectra are considered incorrectly clustered if their peptide labels deviate from the most frequent peptide label in their clusters, with unidentified spectra not considered.

#### Completeness

Completeness measures the fragmentation of spectra corresponding to the same peptide across multiple clusters and is based on the notion of *entropy* in information theory. A clustering result that perfectly satisfies the completeness criterium (value “1”) assigns all PSMs with an identical peptide label to a single cluster. Completeness is computed as one minus the conditional entropy of the cluster distribution given the peptide assignments divided by the maximum reduction in entropy the peptide assignments could provide.^38^

To evaluate scalability, the clustering tools were run on a single, two, and all three test set splits, consisting of 28 million, 56 million, and 84 million MS/MS spectra, respectively. Runtime and memory consumption were measured using the `Unix time` command, using the same hardware set-up as described for training the embedder.

### Clustering peptide annotation

GLEAMS was used to embed all 669 million spectra in the MassIVE-KB dataset and cluster the embeddings. Hierarchical clustering with average linkage and Euclidean distance threshold 0.35 was used, clustering 511 million spectra (76%) with 1.16% incorrectly clustered spectra and 0.837 completeness.

To assign peptide labels to previously unidentified spectra, first peptide annotations were propagated within pure clusters. For 60 million clusters that contained a mixture of unidentified spectra and PSMs with identical peptide labels, the unidentified spectra were assigned the same label, resulting in 82 million new PSMs.

Second, open modification searching was used to process the unidentified spectra. Medoid spectra were extracted from clusters consisting of only unidentified spectra by selecting the spectra with minimum embedded distances to all other cluster members. This resulted in 45 million medoid spectra representing 257 million clustered spectra. The medoid spectra were split into two groups based on cluster size—size two and size greater than two—and exported to two MGF files.

Next, the ANN-SoLo^39,40^ (version 0.3.3) spectral library search engine was used for open modification searching. Search settings included preprocessing the spectra by removing peaks outside the 101*m/z* to 1500*m/z* range and peaks within a 1.5*m/z* window around the precursor *m/z*, precursor mass tolerance 10 ppm for the standard searching step of ANN-SoLo’s built-in cascade search and 500 Da for the open searching step, and fragment mass tolerance 0.05*m/z*. Other settings were kept at their default values. As reference spectral library the MassIVE-KB spectral library was used. Duplicates were removed using SpectraST^41^ (version 5.0 as part of the Trans-Proteomic Pipeline version 5.1.0^42^) by retaining only the best replicate spectrum for each individual peptide ion, and decoy spectra were added in a 1:1 ratio using the shuffle-and-reposition method.^43^ PSMs were filtered at 1% FDR by ANN-SoLo’s built-in subgroup FDR procedure.^44^

ANN-SoLo managed to identify 5.3 million PSMs (12% of previously unidentified cluster medoid spectra). Finally, peptide labels from the ANN-SoLo PSMs were propagated to other cluster members, resulting in 44 million additional PSMs.

## Data availability

The data used to explore the dark proteome have been deposited to the MassIVE repository with the dataset identifier MSV000088598. It consists of MGF files containing the representative medoid spectra from GLEAMS clustering and the associated ANN-SoLo identifications in mzTab format.^27^

All other data supporting the presented analyses have been deposited to the MassIVE repository with the dataset identifier [MSV000088599](https://doi.org/10.25345/C5XW17).

## Code availability

GLEAMS was implemented in Python 3.8. Pyteomics (version 4.3.2)^45^ was used to read MS/MS spectra in the mzML,^22^ mzXML, and MGF formats. spectrum_utils (version 0.3.4)^46^ was used for spectrum preprocessing. Submodular selection was performed using apricot (version 0.4.1).^21^ The neural network code was implemented using the Tensorflow/Keras framework (version 2.2.0).^47^ SciPy (version 1.5.0)^48^ and fastcluster (version 1.1.28)^49^ were used for hierarchical clustering. Additional scientific computing was done using NumPy (version 1.19.0),^50^ Scikit-Learn (version 0.23.1),^51^ Numba (version 0.50.1),^52^ and Pandas (version 1.0.5).^53^ Data analysis and visualization were performed using Jupyter Notebooks,^54^ matplotlib (version 3.3.0),^55^ Seaborn (version 0.11.0),^56^ and UMAP (version 0.4.6).^13^

All code is available as open source under the permissive BSD license at https://github.com/bittremieux/GLEAMS. Code used to analyze the data and to generate the figures presented here is available on GitHub (https://github.com/bittremieux/GLEAMS_notebooks). Permanent archives of the source code and the analysis notebooks are available on Zenodo at doi:10.5281/zenodo.5794613 and doi:10.5281/zenodo.5794616, respectively.

