## Supplementary figures and tables for "A learned embedding for efficient joint analysis of millions of mass spectra"

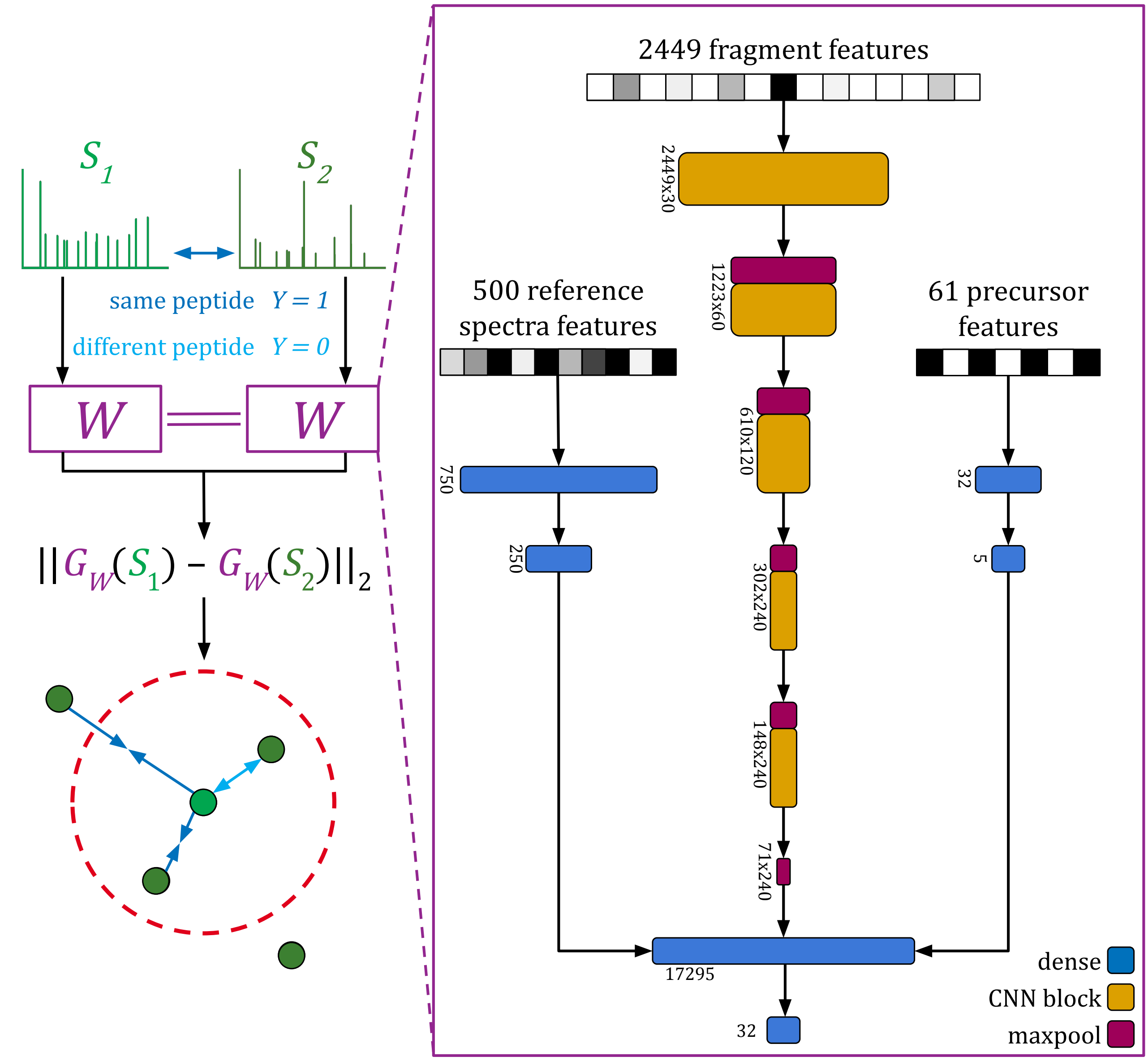

**Supplementary Figure 1:** Each instance of the embedder network in the Siamese neural network separately receives each of three feature types as input. Precursor features are processed through a fully-connected network with two layers of sizes 32 and 5. Binned fragment intensities are processed through five blocks of one-dimensional convolutional layers and max pooling layers. Reference spectra features are processed through a fully-connected network with two layers of sizes 750 and 250. The output of the three subnetworks is concatenated and passed to a final fully-connected layer of size 32.

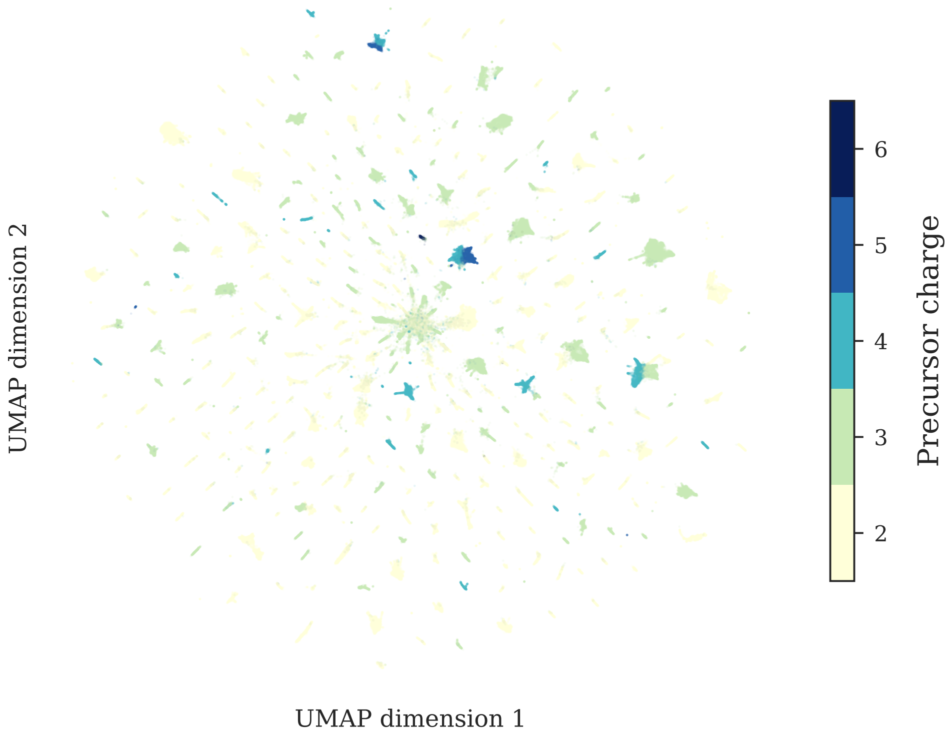

**Supplementary Figure 2:** UMAP projection of 685 337 embeddings from frequently occurring peptides in 10 million randomly selected identified spectra. As indicated by the coloration, precursor charge has a large influence on the location of spectra in the embedded space. Note that the visualization may group peptides with similarities on some dimensions of the 32-dimensional embedding space, but which are nevertheless distinguishable based on their full embeddings.

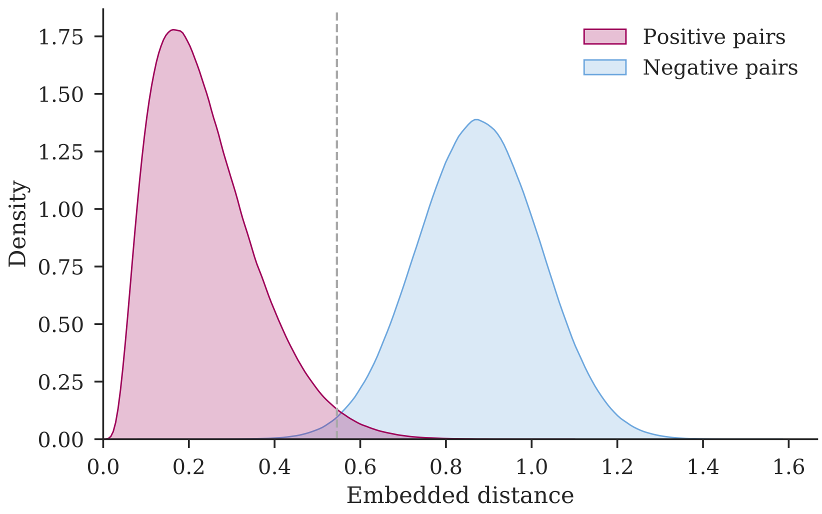

**Supplementary Figure 3:** The false negative rate between positive and negative embedding pairs, for 10 million randomly selected pairs from the test dataset, at distance threshold 0.5455 (grey line), corresponding to 1% false discovery rate, is only 1%, indicating excellent separation between positive and negative pairs.

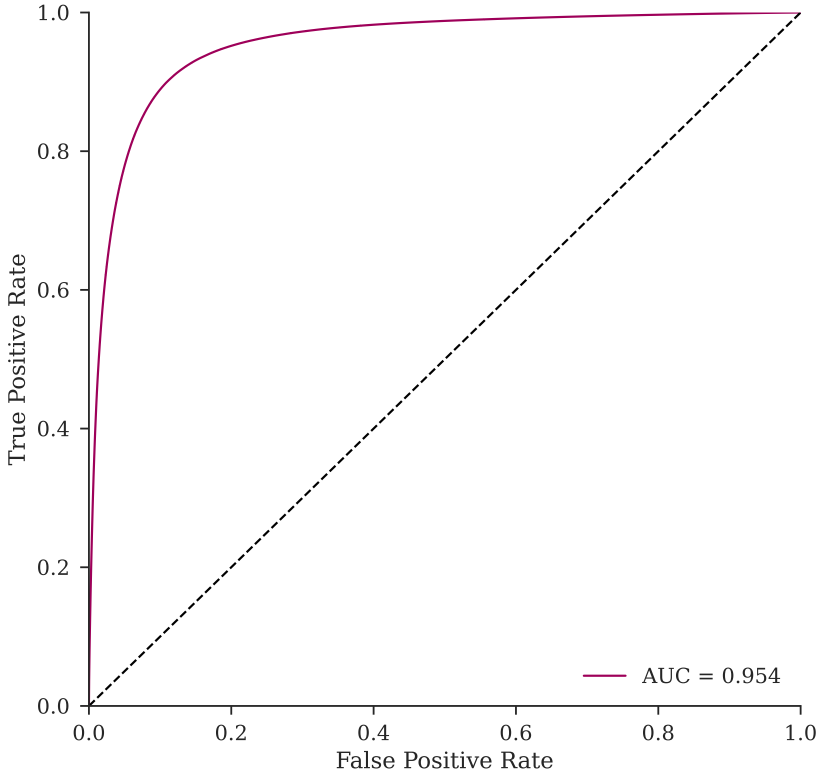

**Supplementary Figure 4:** Receiver operating characteristic curve (ROC) for GLEAMS embeddings corresponding to 7.5 million randomly selected spectrum pairs from an independent phosphoproteomics study.[1] The ROC curve and area under the curve (AUC) show how often a same-peptide spectrum pair had a smaller distance than a different-peptide spectrum pair. Although phosphorylation was not considered as a variable modification during spectrum identification of the MassIVE-KB dataset, and consequently GLEAMS did not encounter any phosphorylated spectra during its training, GLEAMS still managed to correctly assign smaller distances to spectrum pairs corresponding to identical phosphorylated peptide sequences compared to spectra with distinct peptide sequences. This indicates that GLEAMS embeddings are robust toward different types of mass spectrometry data, including unseen modifications.

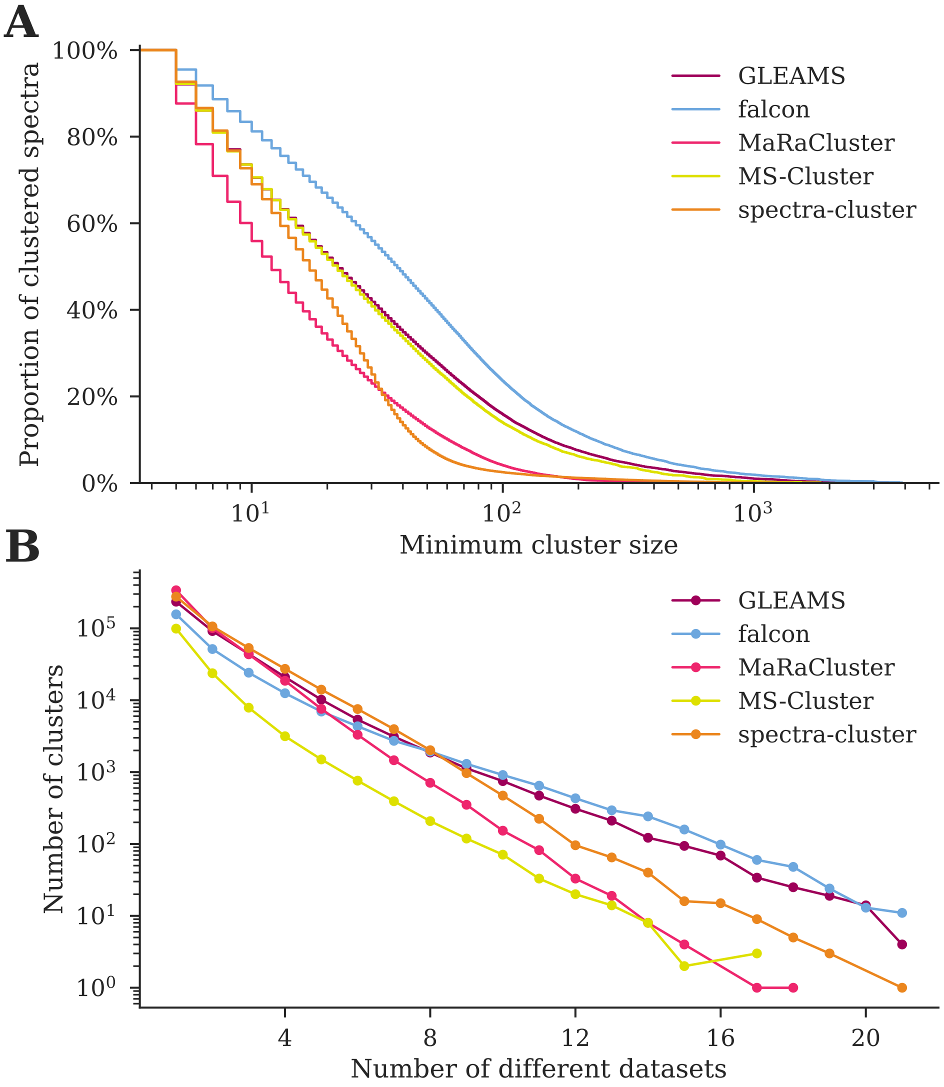

**Supplementary Figure 5:** Clustering result characteristics at approximately 1% incorrectly clustered spectra over three random folds of the test dataset. **(A)** Complementary empirical cumulative distribution of the cluster sizes. GLEAMS produces large clusters, grouping similar spectra into a single cluster rather than splitting them over multiple related clusters. In contrast, MaRaCluster and spectra-cluster predominantly generate clusters containing fewer than 100 spectra. **(B)** The number of datasets that spectra in the test dataset originate from per cluster (24 datasets total). GLEAMS successfully groups related spectra from heterogeneous datasets into single clusters.

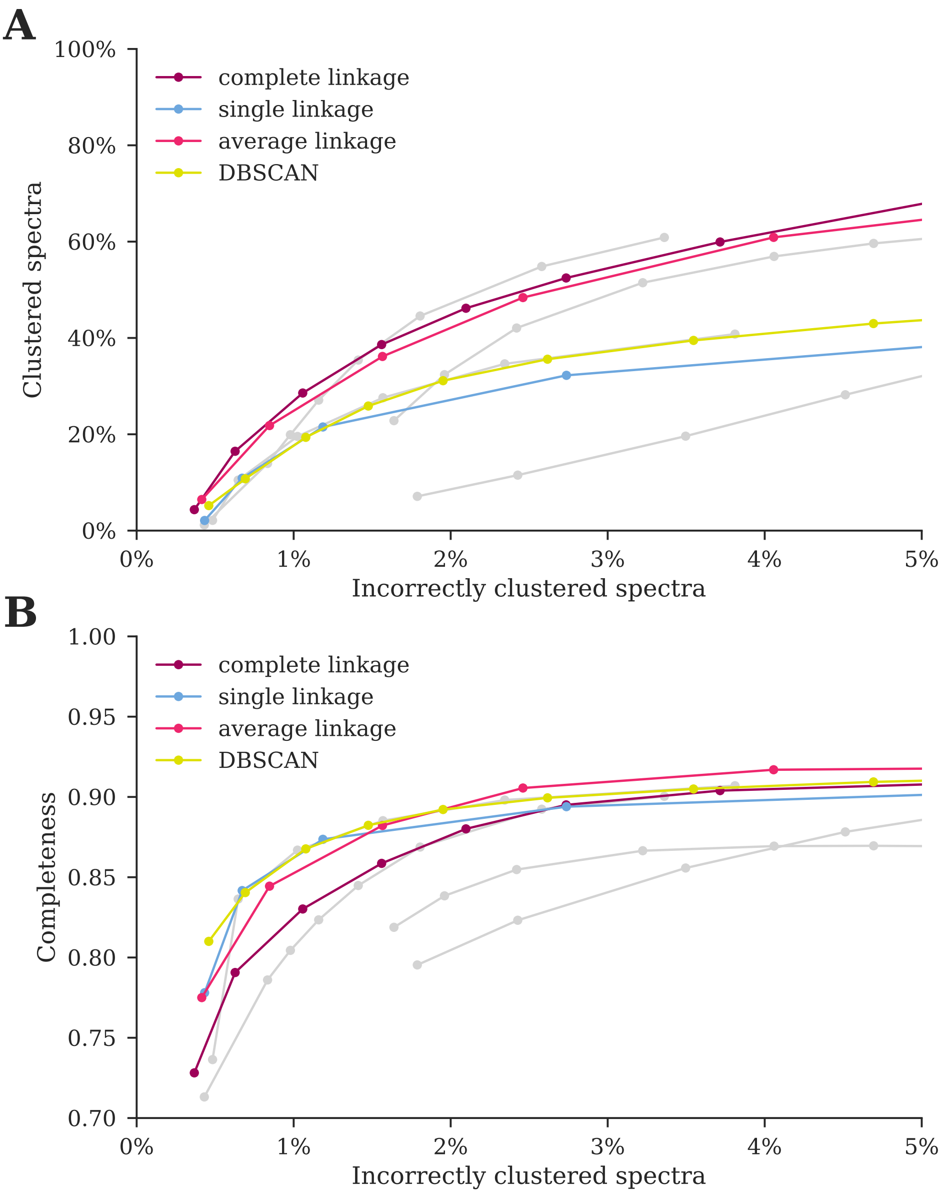

**Supplementary Figure 6:** Average clustering performance over three random folds of the test dataset containing 28 million MS/MS spectra each. The GLEAMS embeddings were clustered using hierarchical clustering with complete linkage, single linkage, or average linkage; or using DBSCAN. The performance of alternative spectrum clustering tools (Figure 1D–E) is shown in gray for reference. **(A)** The number of clustered spectra versus the number of incorrectly clustered spectra per clustering algorithm. Hierarchical clustering using complete linkage or average linkage clusters the highest number of spectra. **(B)** Cluster completeness versus the number of incorrectly clustered spectra per clustering algorithm. GLEAMS consistently achieves optimal completeness, indicating that it successfully groups multiple spectra corresponding to the same peptide in few clusters.

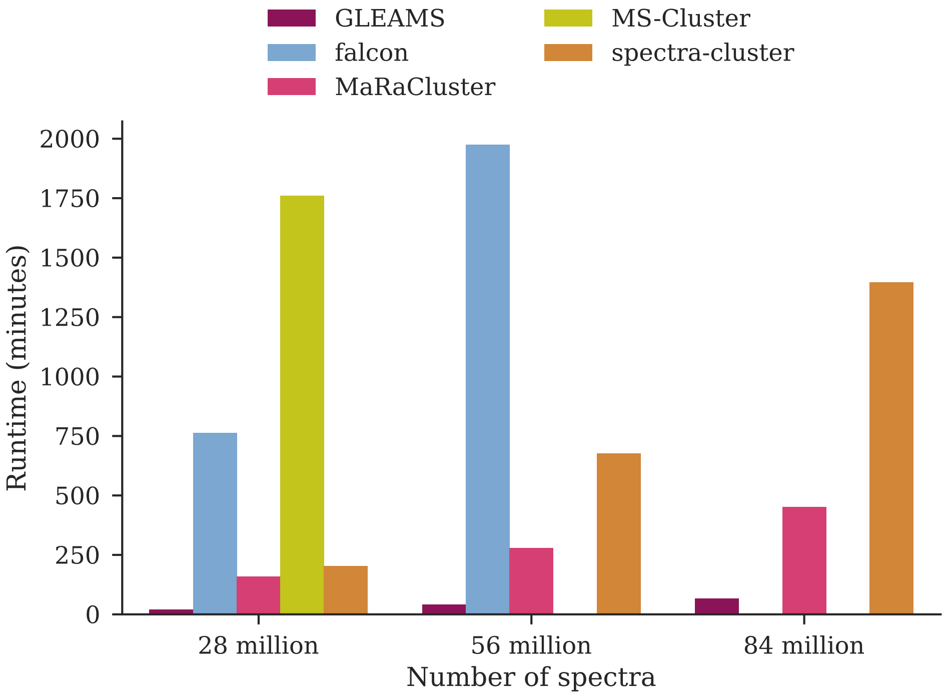

**Supplementary Figure 7:** Scalability of spectrum clustering tools when processing increasingly large data volumes. Three random subsets of the test dataset were combined to form input datasets consisting of 28 million, 56 million, and 84 million spectra. GLEAMS clusters large datasets significantly faster than alternative tools. Evaluations of falcon and MS-Cluster on larger datasets were excluded due to excessive runtimes.

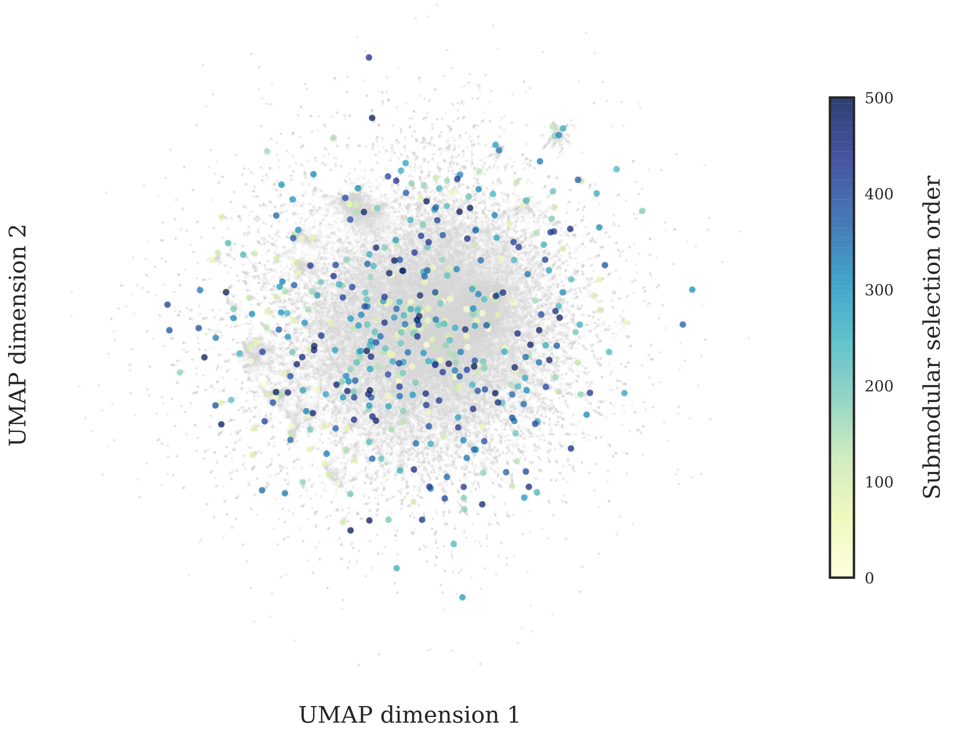

**Supplementary Figure 8:** UMAP visualization of the selected reference spectra. The two-dimensional UMAP visualization was computed from the dot product pairwise similarity matrix between all 200 000 randomly selected spectra from the training data. Submodular selection succeeds in summarizing the full similarity space using 500 representative reference spectra.

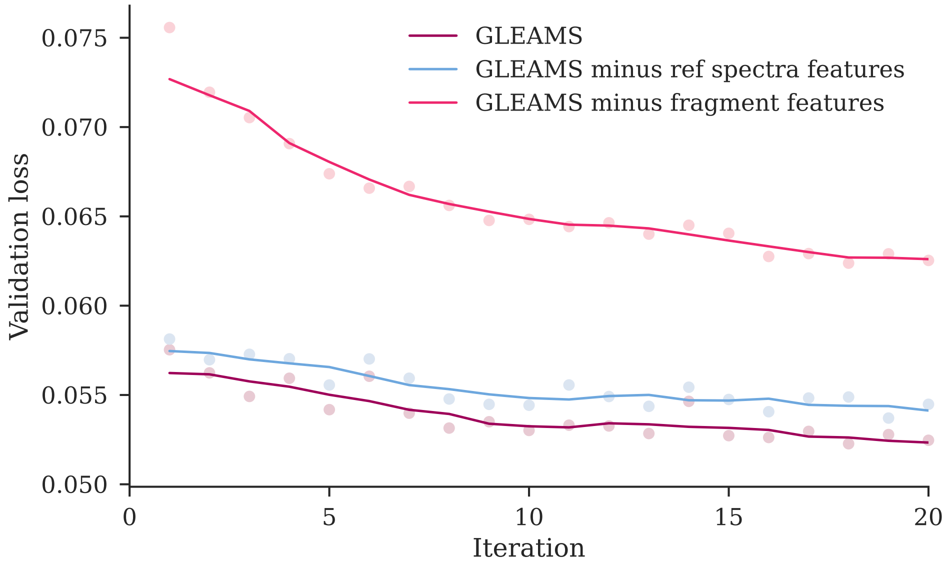

**Supplementary Figure 9:** Ablation testing during training of the GLEAMS Siamese network shows the benefit of the different input feature types. The performance is measured using the validation loss while training for 20 iterations consisting of 40 000 steps with batch size 256. The line indicates the smoothed average validation loss over five consecutive iterations, with the markers showing the individual validation losses at the end of each iteration. Optimal performance is achieved by including all feature types as input to the GLEAMS neural network. Removing either type of feature—reference spectrum features or fragment features—leads to decreased performance (i.e., a higher loss on the validation set).

| Spectrum or peptide property | Embedding dimension | Spearman correlation |
|---|---|---|
| Precursor $m/z$ | 4 | 0.286 |
| | 6 | 0.230 |
| | 7 | 0.218 |
| | 11 | 0.312 |
| | 16 | 0.224 |
| Peptide sequence length | 11 | 0.205 |
| Arginine peptide terminus | 0 | −0.588 |
| | 1 | −0.247 |
| | 4 | −0.212 |
| | 10 | 0.215 |
| | 21 | 0.276 |
| | 22 | −0.219 |
| Lysine peptide terminus | 0 | 0.601 |
| | 1 | 0.237 |
| | 4 | 0.207 |
| | 21 | −0.274 |
| | 22 | 0.214 |

**Supplementary Table 1:** Correlation of individual embedding dimensions with latent properties of the spectra. Spearman correlations above 0.2 and below −0.2 are shown.
